# The transposable elements of the *Drosophila serrata* reference panel

**DOI:** 10.1101/2020.06.11.146431

**Authors:** Zachary Tiedeman, Sarah Signor

## Abstract

Transposable elements are an important element of the complex genomic ecosystem, proving to be both adaptive and deleterious - repressed by the piRNA system and fixed by selection. Transposable element insertion also appears to be bursty – either due to invasion of new transposable elements that are not yet repressed, de-repression due to instability of organismal defense systems, stress, or genetic variation in hosts. Here, we characterize the transposable element landscape in an important model *Drosophila, D. serrata*, and investigate variation in transposable element copy number between genotypes and in the population at large. We find that a subset of transposable elements are clearly related to elements annotated in *D. melanogaster* and *D. simulans*, suggesting they spread between species more recently than other transposable elements. We also find that transposable elements do proliferate in particular genotypes, and that often if an individual is host to a proliferating transposable element, it is host to more than one proliferating transposable element. In addition, if a transposable element is active in a genotype, it is often active in more than one genotype. This suggests that there is an interaction between the host and the transposable element, such as a permissive genetic background and the presence of potentially active transposable element copies. In natural populations an active transposable element and a permissive background would not be held in association as in inbred lines, suggesting the magnitude of the burst would be much lower. Yet many of the inbred lines have actively proliferating transposable elements suggesting this is an important mechanism by which transposable elements maintain themselves in populations.

## Introduction

Transposable elements (TEs) are short sequences of DNA that multiply within genomes despite potential deleterious impacts to the host (McClintock 1950). TEs are widespread across the tree of life, often making up a significant portion of the genome (Piegu et al. 2006; Schnable et al. 2009; Lee and Langley 2012). TEs also impose a severe mutational load on their hosts by producing insertions that disrupt functional sequences and mediate ectopic recombination (McGinnis et al. 1983; Levis et al. 1984; Lim 1988). TEs can spread through horizontal transfer between non-hybridizing species, allowing them to colonize new host genomes (Kidwell 1983; Kofler et al. 2015; Peccoud et al. 2017). For example, the spread of the *P*-element was documented in *D. melanogaster* from *D. willistoni* in the 1950’s, and its subsequent spread into *D. simulans* around 2010 (Daniels et al. 1990b; Kofler et al. 2015).

TEs have also been implicated in adaptation. In *Drosophila*, insertion of TEs has been linked to resistance to pesticides and viral infection (Wilson 1993; Daborn et al. 2002; Aminetzach et al. 2005; Magwire et al. 2011; Mateo et al. 2014). In ants and *Capsella rubella*, TEs provide genetic diversity in invading populations which are generally depleted of variation, facilitating adaptation to novel environments (Schrader et al. 2019; Niu et al. 2019). In fission yeast, TE activity was increased in response to stress and TE insertions were associated with stress response genes, supporting the supposition that TEs provide a system to modify the genome in response to stress (Esnault et al. 2019). There is also evidence from vertebrates that TEs provide the raw material for assembling new protein architectures through capture of their transposase domains (Cosby et al. 2020). In summary there is extensive evidence that TEs provide genetic material for adaptation through a variety of mechanisms.

Despite the evidence for an adaptive role for TEs, most TE insertions are thought to be deleterious, and the host has a dedicated defense mechanism termed the PIWI-interacting (piRNA) system. piRNAs bind to PIWI-clade proteins, such as *Argonaute 3* in *D. melanogaster*, and suppress transposon activity transcriptionally and post-transcriptionally (Brennecke et al. 2007). The majority of these piRNAs originate from genomic regions which are enriched for TE fragments, termed piRNA clusters (Brennecke et al. 2007; Malone et al. 2009). These piRNA clusters are large, and there is at some evidence that insertion of a TE into a piRNA cluster is enough to initiate piRNA mediated silencing of the TE (Josse et al. 2007; Zanni et al. 2013). Therefore a newly invading TE would proliferate in the host until a copy jumps into a piRNA cluster, which then triggers piRNA silencing of the TE (Bergman et al. 2006; Malone and Hannon 2009; Zanni et al. 2013; Goriaux et al. 2014; Yamanaka et al. 2014; Ozata et al. 2019). These piRNA clusters are preferentially located in heterochromatic regions and usually have low recombination rates (Brennecke et al. 2007). This reduces the efficacy of purifying selection and may serve as ‘safe harbors’ for TEs to accumulate and develop into piRNA clusters (Brennecke et al. 2007; Kofler 2019; Zhang and Kelleher 2019). piRNA evolution is thought to be rapid enough that adaptation to a novel TE could occur within the lifetime of an individual (Khurana et al. 2011). The transposition rate of TEs is also controlled by other mechanisms, including regulation of promotor activity, chromatin structure, and splicing (Guerreiro 2019). In some cases the mechanism is unknown, such as the accumulation of *copia* in the genomes of inbred *D. melanogaster*, suggesting that there is still more to know about the regulation of TE copy number (Pasyukova 2004).

This apparent contradiction, between the existence of a dedicated repression machinery, and an apparent important role for TEs in adaptation, also complicates inferences about the tempo and mode of TE transposition. For a long time, active TEs variants were thought to be rare in natural populations (Kaplan et al. 1985; Ronsseray et al. 1991; Brookfield 1991; 1996; Nuzhdin et al. 1997). Or, it was not active TEs which are rare but individuals with ‘permissive’ genetic backgrounds, such that TEs would remain inactive until encountering a permissive genetic background and then proliferate (Nuzhdin 2000). Either way, these models assumed a transposition – selection balance such that TEs proliferated at approximately the rate that they were removed by selection. Since then, TEs have been observed to undergo bursts of activity, which could occur for multiple reasons such as colonization, hybridization, and stress (Vieira et al. 1999; Romero-Soriano and Garcia Guerreiro 2016; Guerreiro 2019). These bursts are documented in *Drosophila*, rice, fish, and other systems (Vieira et al. 1999; Piegu et al. 2006; de Boer et al. 2007; Bourgeois and Boissinot 2019; Signor 2020). In most cases, transposition bursts in *Drosophila* include few individuals and TEs (Biémont et al. 1987; 1990; Nuzhdin et al. 1997; Yang et al. 2006). The underlying explanation for this burstiness is unclear, including the potential role of burstiness in adaptation. Bursts of transposition would be expected upon invasion of a new TE, prior to silencing by the piRNA system, however TEs also appear to become reactivated in response to stress, or potentially variation in the host suppression system.

Recently an inbred panel of 110 genotypes was created for *D. serrata*, a member of the *montium* subgroup (Reddiex et al. 2018). The *montium* group contains 98 species and represents a significant fraction of known *Drosophila* species (Lemeunier et al. 1986; Reddiex et al. 2018). The *D. serrata* panel was sampled from a single large population within its endemic distribution in Australia (Reddiex et al. 2018). *D. serrata* is a model system for understanding latitudinal clines and the evolution of species boundaries (Blows 1993; Jenkins and Hoffmann 1999; Hallas et al. 2002; Hoffmann and Shirriffs 2002; Liefting et al. 2009). While the development of a panel represents a new opportunity for genomic investigation in the group, such as GWAS, very little work has been done understanding the landscape of repetitive elements in this group. For example, *D. serrata* was found to contain a domesticated *P*-element, though no evidence of active *P*-elements was noted (Nouaud and Anxolabéhère 1997; Nouaud et al. 1999). Screens for the presence of the *Drosophila hobo* element in the *montium* group were mixed, and inconclusive for *D. serrata* (Daniels et al. 1990a). *copia* and *412* were not detected in *D. serrata*, though the DNA transposon *Bari-1* was (Biémont and Cizeron 1999), and evidence for the presence of the *mariner* element is equivocal (Maruyama and Hartl 1991; Brunet et al. 1994). Here we will characterize the TE landscape in the *Drosophila serrata* Genetic Reference panel. This will have two goals: 1) To understand the TE content of *D. serrata* and its relationship to existing TE annotations 2) To understand variability in TE content between individuals in the population and how this relates to the tempo and mode of TE movement. This will provide the groundwork for understanding the role of TEs in evolution in *D. serrata*, as well as provide another investigation into the proliferation of TEs in individual genetic backgrounds.

## Methods

### Fly lines and data

110 genotypes of *D. serrata* were collected from a wild population in Brisbane Australia in 2011 and inbred for 20 generations (Reddiex et al. 2018). The libraries were sequenced using 100 bp paired-end reads on an Illumina Hi-seq 2000. The raw reads were downloaded from NCBI SRA PRJNA410238. 104 genotypes were used for analysis. 4 genotypes were excluded based on unusually high relatedness, as described in (Reddiex et al. 2018), while the remaining 2 genotypes were excluded based on library quality issues.

### Classification of TEs

TEs are a diverse group, and the taxonomy of TEs is contentious and still developing (Wicker et al. 2007; Kapitonov and Jurka 2008; Platt et al. 2016). Here, we will rely only on broad classifications in Class I and Class II elements, including *Helitrons* and miniature inverted-repeat TEs (MITES). Class I elements are retrotransposons which use an RNA intermediate in their ‘copy and paste’ transposition. Class I can be divided into long terminal repeat (LTR) and those that lack LTRs (SINEs and LINEs) (Okada et al. 1997; Havecker et al. 2004; Wicker et al. 2007; Kramerov and Vassetzky 2011; 2019). However, here we will only focus on LTRs in Class I, as benchmarking of software designed to detect non-LTRs is unreliable (Ou et al. 2019). Within the LTRs, there are two major superfamilies – *copia* and *gypsy* – which have distinct terminal sequences (Marlor et al. 1986). Class II elements are known as DNA transposons, or terminal inverted repeat transposons (TIR), and use DNA intermediates in a ‘cut and paste’ mechanism of transposition (McClintock 1984). Among the TIRs are also non-autonomous small DNA transposons such as miniature inverted-repeat TEs (MITES) (Fattash et al. 2013; Makalowski et al. 2019). These can belong to any of the described TIR superfamilies, but they lack coding potential and rely on other autonomous DNA transposons for transposition. Lastly, the *Helitron* TIR was discovered in 2001 and has a different mechanism of transposition, referred to as a rolling circle, which frequently captures nearby genes or portions of them in the process (Kapitonov and Jurka 2001; Kapitonov and Jurka 2007).

### Mapping and copy number estimation

The *D. serrata* 1.0 assembly available from the Chenoweth lab was used for genomic mapping and TE identification (http://www.chenowethlab.org/resources.html) (Allen et al. 2017). The TE library was constructed using the Extensive de-novo TE Annotator pipeline (EDTA) (Ou et al. 2019). This pipeline is intended to create a high quality non-redundant TE library based off of a reference genome. Reads from the *D. serrata* reference panel were mapped to the genome and the TE library using bwa mem version 0.7.15 (Figure 1; Li 2015). Bam files were sorted and indexed with samtools v.1.9 and optical duplicates were removed using picard MarkDuplicates (http://picard.sourceforge.net) (Li et al. 2009; McKenna et al. 2010). Reads with a mapping quality of below 15 were removed (this removes reads which map equally well to more than one location). Using read coverage to determine copy number has been compared to other methods and is neither permissive nor conservative (Srivastav and Kelleher 2017). TE copy number was estimated using the average counts of reads mapping to the TE sequences and the genome with bedtools counts (Quinlan and Hall 2010; Hill et al. 2015). Then, copy number of the TEs could be normalized using the average counts from a 7 MB contig from *D. serrata* which corresponds to a portion of *D. melanogaster* 3L. This is one of the largest contigs in the *D. serrata* assembly.

**Figure 1:**
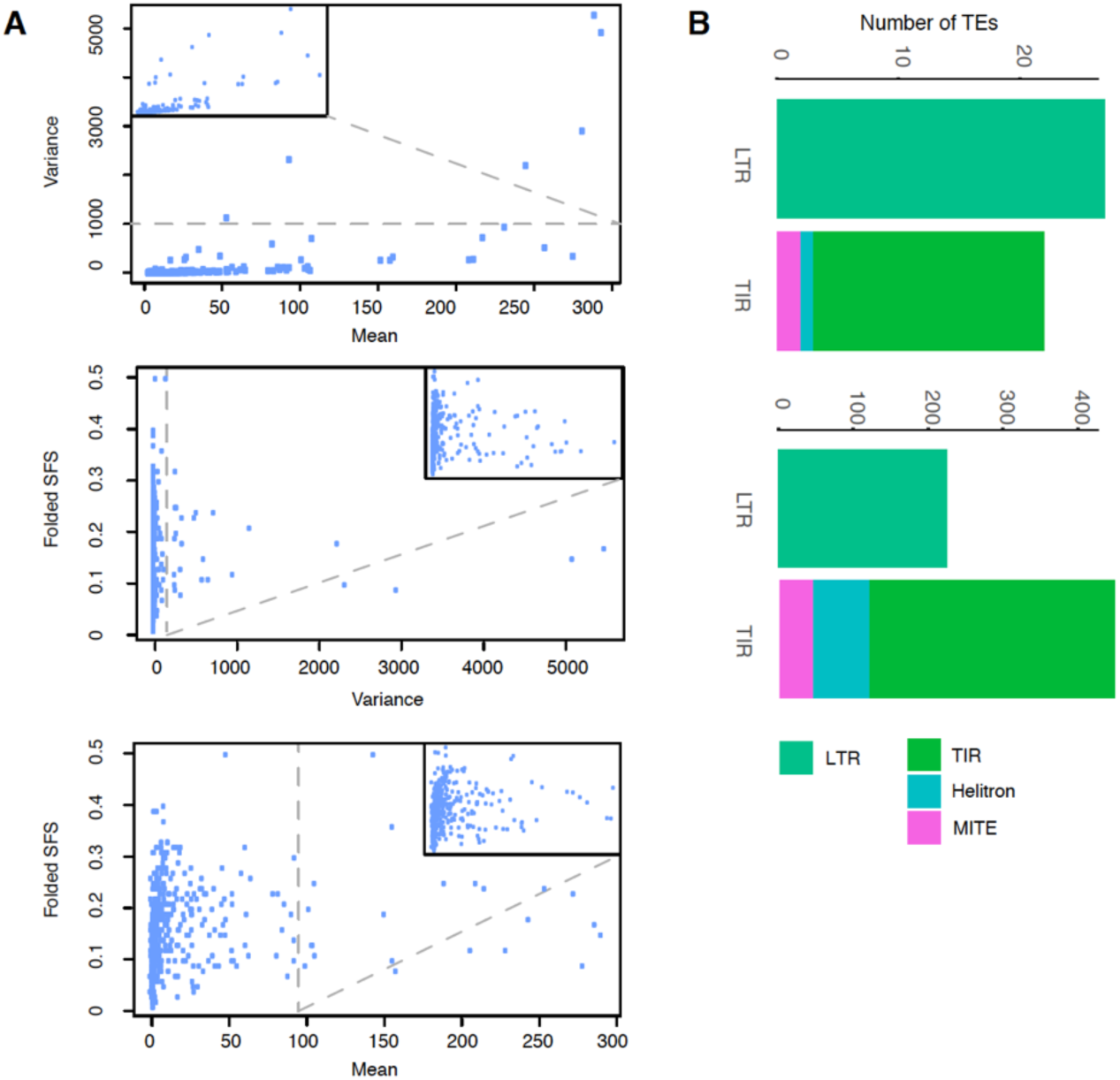
A. Mean and variance of copy number as well as folded SFS in transposable elements of Australian *D. serrata*. On the top, the majority of identified transposable elements have low mean copy number and variance within the population. The top three most variable elements are excluded from all graphs so as not to compress the y-axis. B. The 52 most variable transposable elements (top) with variance greater than 50 versus all of the transposable elements (bottom). Overall the number of LTRs represent fewer of the total identified TEs, however of those that are highly variable they are equally likely to be LTRs or TIRs.

### SNPs and summary statistics

We called SNPs within the TEs using GATK Haplotypecaller (McKenna et al. 2010). SNPs were not filtered for missing calls given that not all individuals will share insertions. However, they were filtered for a coverage of at least 4 reads to be called in an individual. The site frequency spectrum (SFS) of SNPs in the TEs was estimated with VCFtools as the frequency of each SNP in the population, and then the frequency of the SNP frequencies was estimated in R (Danecek et al. 2011). The SFS was then folded in R, replacing any frequency *i* over 0.5 with 1-*i*. This was done because we could not determine the derived allele.

### Relationship to TEs in the EMBL TE library

The TE library from *D. serrata* was compared to TEs from the EMBL library using DFAM (Hubley et al. 2016). Hits were required to have a bit score of greater than 350. Multiple hits to the same TE were considered as a single hit, and if more than one EMBL TE was listed the best bit score was retained. In general there were no TEs from the *D. serrata* library that had similar bit scores between different EMBL TEs.

### Relationship between TEs annotated by EDTA

Potentially related TEs from the EDTA library were identified using ncbi blastn 2.8, with the minimum criteria being an alignment of greater than 400 bp for LTRs and TIRs, and 200 bp for MITEs (Camacho et al. 2009). The sequences were aligned and oriented using the R package DECIPHER (Wright 2016). The fasta alignments were converted to nexus format, and indels were coded as binary characters, using the perl script 2matrix (Salinas and Little 2014). Trees were made if there were four or more related TEs using MrBayes 3.2.7 (Ronquist et al. 2012). The trees were built using a GTR substitution model and gamma distributed rate variation across sites. The markov chain monte carlo chains were run until the standard deviation of split frequencies was below .01. The consensus trees were generated using sumt conformat=simple. The resulting trees were displayed with the R package ape (Paradis et al. 2004).

## Results

### EDTA

EDTA identified 676 TEs in the *D. serrata* reference genome. The sequences of these TEs are available at https://github.com/signor-molevol/serrata_transposable. The classification of the TEs into superfamilies is broadly correct, and in many cases there is no clear relationship to an existing TE. However, some errors are evident, for example, element *444* is classified as *copia*, but aligns quite well with the *297* element in *D. melanogaster*, which is a member of the *17*.*6* clade/*gypsy* superfamily. In addition, some unknown elements such as *69* align well with existing *D. melanogaster* annotations, in this case *17*.*6*. In all 6 elements that were classified as unknown or *copia* align well with members of the *gypsy* superfamily from *D. melanogaster*. Therefore classification below the superfamily level is generally ambiguous, though MITEs, *Helitrons*, and TIRs are distinguishable. This may be due to deletion of canonical sequences, nested insertions, or other ambiguities of TEs.

### Population frequency of TEs

An average of 17% of reads from individual *D. serrata* lines mapped to TE sequences. The average number of TEs per genome in this population of *D. serrata* is 19,909, however almost 50% of that total (9,036) are from a single repetitive uncharacterized sequence (Supplemental File 1). This sequence is classified as an LTR, though it does not share sequence similarity to other well characterized LTRs. This element shares a 36 bp segment with *D. melanogaster INE-1*, and may be misclassified given that *INE-1*s are generally very abundant. The next closest in copy number is a TIR with 541 copies, thus this is a significant outlier. 6 TEs identified in the reference are likely not present in this population. 2 of these are present as partial copies in a subset of individuals. Overall among the elements identified by EDTA approximately twice as many are TIRs compared to LTRs (Figure 1). However, the majority of the identified TEs have low copy number and variance. 390 of the identified elements have an average copy number of less than 3, and all but 2 of those have a variance of less than 1 (Figure 1, Supplemental file 1; the other two have variances of 3 and 4). Of those remaining, 148 have a variance of 3 or less (Figure 1). Therefore the vast majority of TEs in this population vary little in copy number (Figure 1). However, among those that do vary considerably, 52 elements have a variance in the population greater than 50. These represent a very different subset of TEs than those identified overall – an approximately equal number are LTRs and TIRs (Figure 1). This suggests that LTRs are more active in the population, which is consistent with other work on transposable elements that found that LTR insertions were generally of more recent origin than TIRs (Kofler et al. 2015).

### Folded SFS

Overall ∼6% of SNPs in TEs have a frequency of higher than 60% in this population. Rather than determine the ancestral state by adding an outgroup, we chose to fold the allele frequency spectrum. Overall the folded SFS is low (average of 0.13), however SNPs that are not in TEs also have excess of low frequency variants genome-wide according to genome-wide Tajima’s *D* (Reddiex et al. 2018). There is not a clear relationship between the folded SFS and mean/variance (Figure 1A). A folded SFS of greater than 0.2 is associated with lower copy number and variance overall (Figure 1A). 16 elements have a folded SFS of 0.3 or higher, and the majority of these have low variance in copy number (2 or less) suggesting that they are not active and have been diverging. There are two obvious types of TEs that have likely been spreading recently in the population as a whole – those with no SNPs or low SFS, and with high variance and/or copy number. 5 TEs have no SNPs and therefore no folded SFS can be calculated (*217, 411, 610, 624, 638*, Table 1, Figure 2). This includes *610*, a *gypsy* element which aligns to the internal sequence of *Dsim\ninja* suggesting it is distantly related, but more recently moved into *D. serrata* than TEs with no obvious relatives in related species. As with the population of TEs as a whole, most TEs with a low SFS (lower than .05) also have low copy number (less than 3) and low variance (less than 1). Of these 71 elements, 4 are an exception and have a low SFS, higher copy number, and higher variance. This includes a MITE element and three TIRs. A lack of SNPs, or low SFS, along with higher copy number and variance, suggests that the TEs have been spreading recently in the population (Table 1).

**Figure 2:**
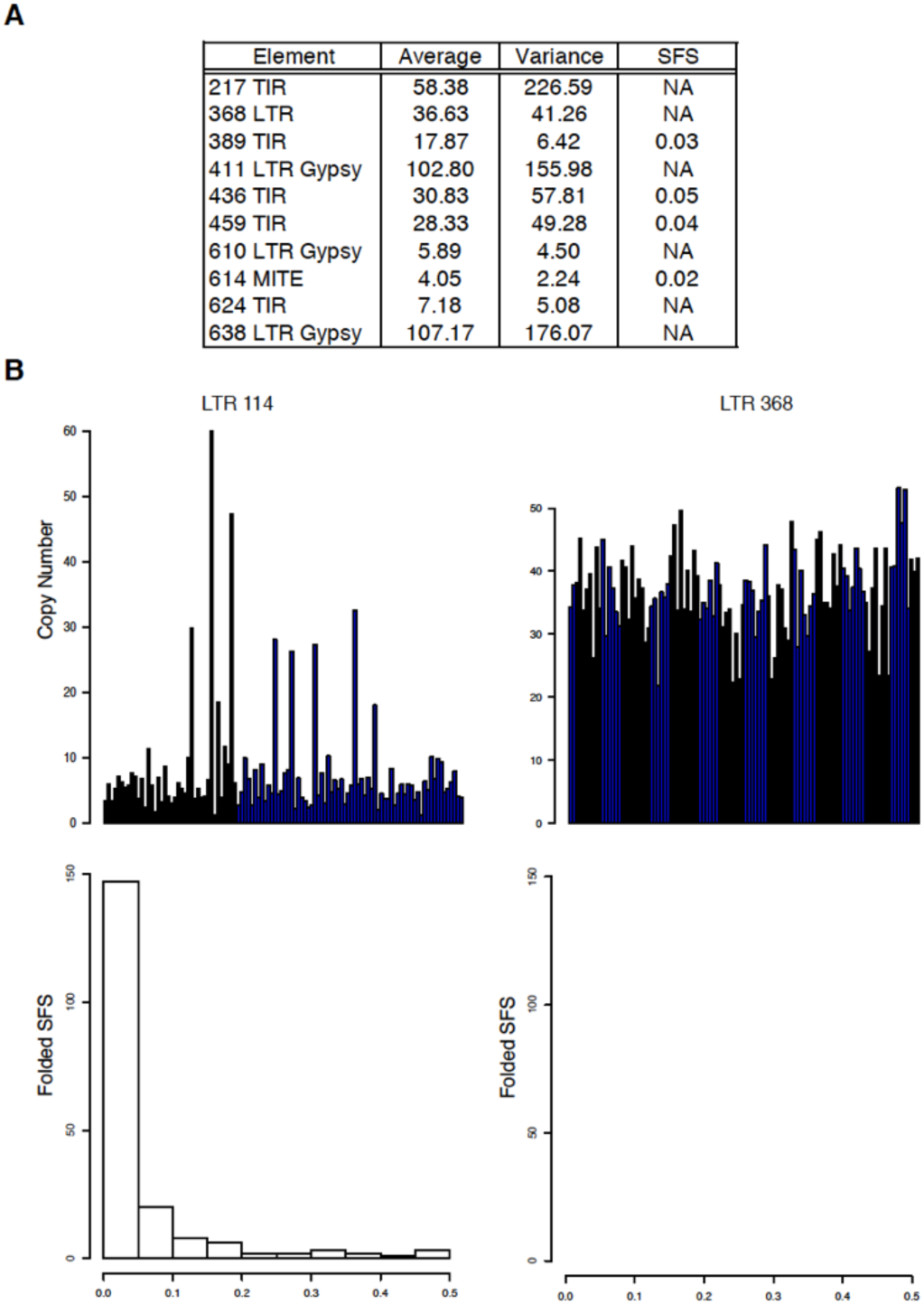
A. A list of transposable elements with variance of at least 2 and either no SNPs or and SFS of 0.05 or less. B. Variation in copy number in the population for two transposable elements (top) and the folded SFS (bottom).

However, TEs may also have been in the population for long enough to accumulate SNPs, but experience bursts of activity. Among the 52 TEs with a variance higher than 50, 17 TEs have an SFS of greater than 0.2 suggesting they have been in the population for longer but have had active transposition of divergent copies (Supplemental Table 1). This group consists of 15 LTRs, one MITE, and one TIR. This could be due to the presence of older copies accumulating SNPs compared to younger active copies, or divergence between different active copies in the population, or both.

### Outliers in individual genotypes

TEs tend to proliferate in particular inbred genotypes. Out of 104 genotypes, 73 have no TEs with a number of insertions that classify them as outliers. 12 genotypes contain a single TE with a copy number that is considered an outlier, and the remaining 19 contain 2 or more outliers. This includes 2 genotypes with 13 and 8 TEs with a copy number that is considered an outlier. This also tends to group by TE, as only 36 TEs have at least 1 genotype in which they are an outlier, however for 18 of these this is only in 1 genotype. For 5 genotypes, 5 TEs are shared as being outliers, with an additional 2 genotypes which share outliers for 4 of the 5. Many of these outliers are large, for example for element *512* the majority of the population has 20-30 copies, while a single individual has > 200 (Figure 3).

**Figure 3:**
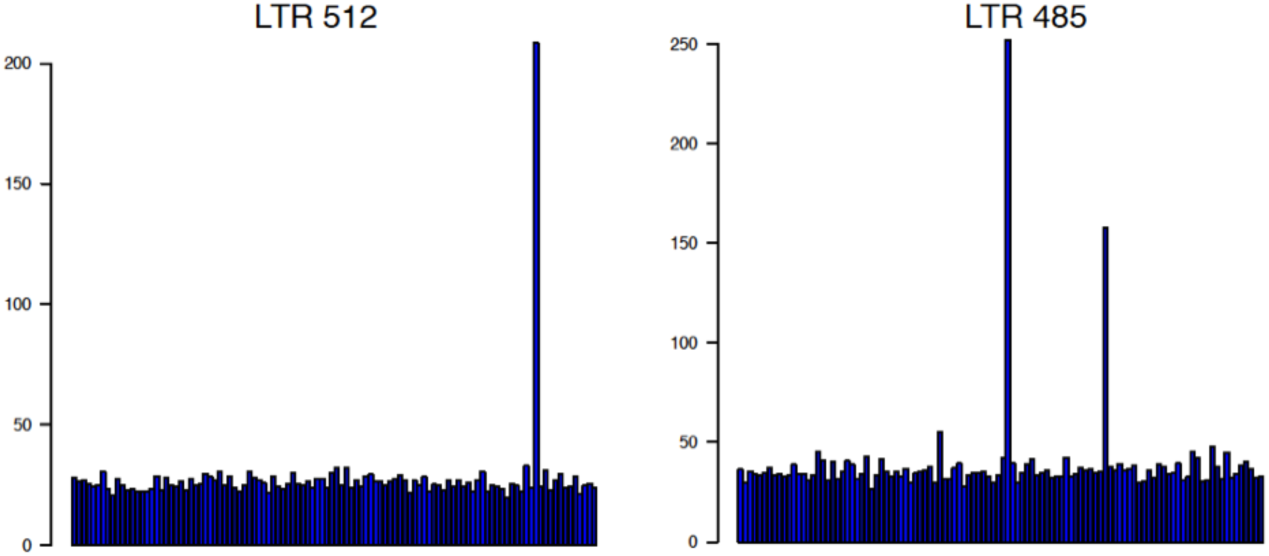
An example of genotypes with an accumulation of transposable elements. In both panels the population aver is 20-30, while individual genotypes have in excess of 150 copies.

### Relationship to existing TE annotations

123 of the 676 identified elements have a well-supported relationship to existing DFAM TE annotations (Figure 4). This includes, for example, 27 elements that are related to the *D. melanogaster Max-Element* and 10 elements that are related to the*D. simulans ninja* element. One of these is also among the most variable TEs (variance greater than 50), and is most closely related to the *Circe* element (*Osvaldo* family). These are likely to be TEs that moved between species more recently, and they are almost exclusively LTRs. The exception being two TIRs from the *hobo* family, one *Helitron* from *D. melanogaster*, and two *Helitrons* most closely related to elements from *Heliconius*. No evidence of *P*-elements were found in the population of identified TEs. In addition, *jockey* elements (non-LTR retrotransposons) are not intended to be identified as a part of this pipeline but do appear to be the identity of two transposons. The overall phylogeny of the TEs is not what we wish to emphasize here, as the structure of TE classification changes frequently (for example whether something is a clade or a family, etc.). In *Drosophila* there is evidence that *gypsy* elements are infectious, as they can be transferred among strains through exposure or microinjection (Song et al. 1994; Kim et al. 1994). This makes them more likely to spread between species.

**Figure 4:**
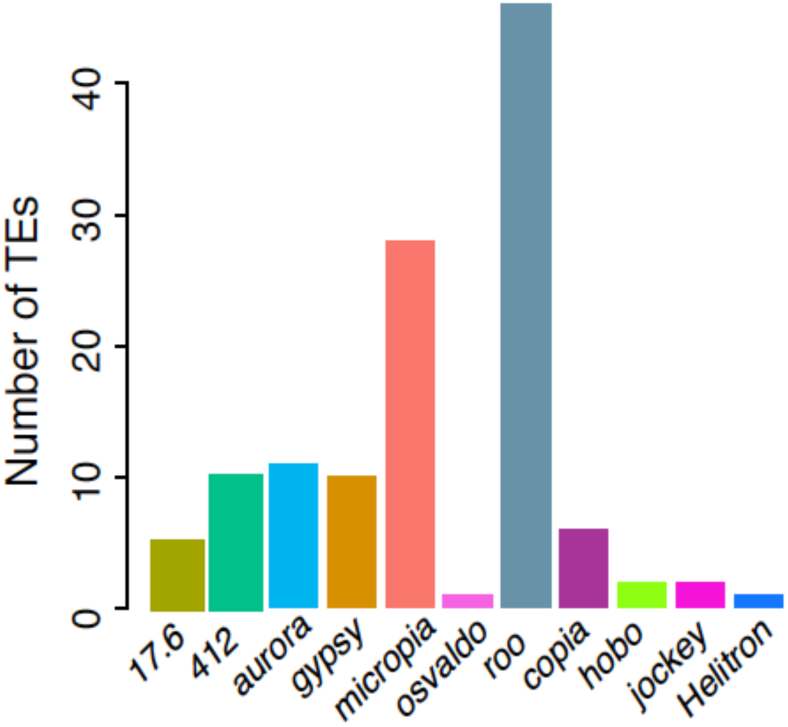
The classification of transposable elements into clades which could be tied to annotated *D. melanogaster* elements. The two *Helitron* elements potentially related to those from *Heliconius* are not included.

**Figure 5:**
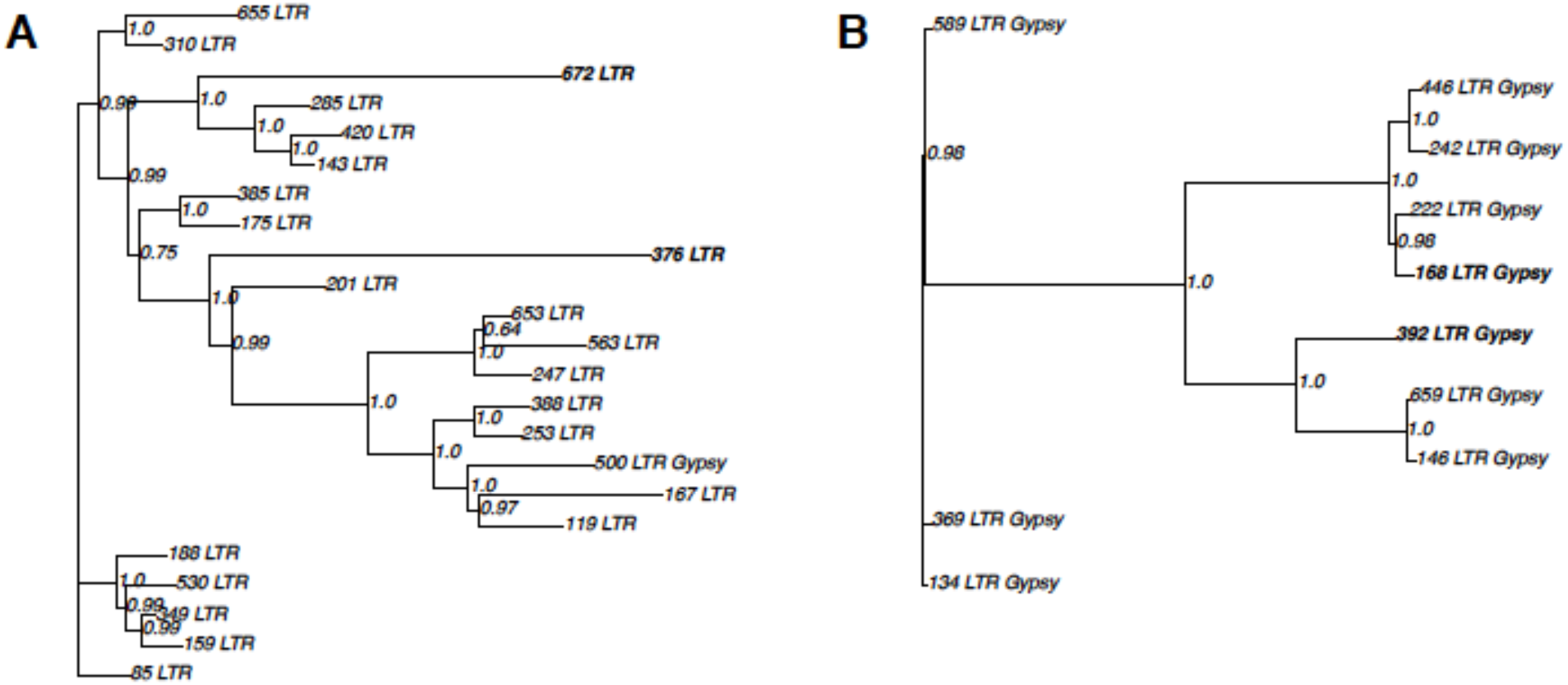
Relatedness between groups of TEs annotated by EDTA. Posterior probabilities of each division is shown, however branch lengths are not meaningful given that TEs do not follow a standard substitution model. This does not represent only the degradation of old copies of TEs, different members of the TE families continue to likely be active, shown in bold.

### Relationship between TEs annotated by EDTA

40 groups of 170 TEs annotated by EDTA are clearly related to one another (Supplemental File 3). For example, 8 TEs (*52, 60, 276, 346, 367, 424, 539, 601*) share sequence similarity for the entirety of their length, but are separated by 39 deletions spread across the TE. In the largest related group of TEs, 23, most of the versions of this TE have low copy number and variance (Figure 4A, average copy number 2.3, average variance < 1). However, two members of the group are likely still active and have relatively high copy number and variance (*376* and *672*, copy number 27, 79; variance 10, 102). Note that the more active TEs do not group together (shown in bold), however because TEs cannot be assumed to follow a standard substitution model the branch lengths are not meaningful (Figure 4A). In another case, 3 members of the group are more distantly related, while 7 members are more closely related and form 2 clear groups of origin (Figure 4B). Yet again, those which are active in the population, as evidenced by higher copy number and variance, are not the most closely related (Figure 4B, shown in bold)

The distance between TEs cannot be understand in the same way as genomic DNA, however in some cases the TEs are clearly related by a simple tree of SNPs and indels (as above), and in others it is more complicated. For example, in a group of 4 TEs (*237, 225, 358, 468*) the long terminal sequence of the TEs is nearly identical and allies them with *Max* elements, however member *468* appears to also carry an unshared insertion of a portion of a *297* element, among other complicated indels. *468* has a copy number of 5, indicating it continued to be active while carrying a portion of a *297* element, though it has low variance between individuals. Interestingly, in another case these relationships appear to describe the origin of 3 MITEs (*399, 405, 472*) from a parental TIR (*660*). The parental TIR has a high copy number in the population, with an average of 286 and a variance of more than 6,000. This is not intended to be an exhaustive accounting of relationships between these TEs, for example at some point all members of the *roo* clade shared an ancestor. Rather, this is intended to describe recent divergence between members of a group within this species.

## Discussion

There is an abundance of evidence from inbred lines that genotypes can vary considerably in copy number. The question remains – is it due to differences in the permissiveness of the genetic background, or inheritance of active TEs that segregate at low frequency in the population? In the former scenario, genes segregating in natural populations modify transcription and the rate of transposition of specific TEs, including polymorphisms in genes such as *Argonaute 3* and variation in the integration of TEs into piRNA clusters (Birchler et al. 1989; Pélisson et al. 1994; Csink et al. 1994; Lee and Langley 2010; 2012; Zhang and Kelleher 2019). Indeed, variation in the integration of TEs into piRNA clusters appears to be quite common, as Zhang and Kelleher (2019) documented 80 unique independent insertions of *P-elements* into piRNA clusters in the *Drosophila* Genetic Reference Panel (Mackay et al. 2012). If laboratory lines differ in these alleles, this can cause between line variability in transposition rates. In the latter scenario, different lines may have inherited copies of TEs with differences in the propensity to transpose (Ronsseray et al. 1991; Kim et al. 1994; Nuzhdin et al. 1997; Nuzhdin 2000).

While we cannot measure the likelihood of individual genotypes inheriting multiple active copies of TEs while fellow members of the population inherit none, the fact that multiple TEs are proliferating in individual genotypes supports the idea that these individuals have polymorphisms in genes or other repressive structures that are more permissive to TE transposition. Were the genotypes with clear TE proliferation different for every TE family this would not support either scenario, however it does seem more likely that these genotypes have a polymorphism which fails to repress more than one type of TE, rather than that they preferentially inherited multiple active copies. We cannot at this time directly look for polymorphisms in repressive genes or complexes. Currently we are unable to establish clear homologs of the *D. melanogaster* genes known to affect piRNA silencing in *D. serrata*, but as the *D. serrata* assembly improves this may be possible. In addition, the methods developed recently be Zhang and Kelleher (2019) to measure differences in piRNA cluster integration using small RNA libraries shows promise for determining whether we can detect polymorphisms in these individual genotypes for repressive alleles.

However, the fact that the TEs which are proliferating do not appear to be a unique population suggests that there is interaction between potentially active TEs and genetic background – not all TEs are potentially active in all potentially permissive backgrounds. This suggests that the transposition rate of TEs in natural populations will be complex, depending upon differences in the inherited TE population and variation in the host genome. There is already a lot of evidence that there are multiple pathways and factors that control transposition in *Drosophila*. For example, in *D. melanogaster* strain *iso-1* the piRNA pathway produces normal *hobo* and *I*-element specific piRNAs, yet there is a high level of *hobo* and *I*-element transposition (Zakharenko et al. 2007; Shpiz et al. 2014). In *D. simulans*, there is large amounts of variation in piRNA pathway genes (Fablet et al. 2014). Therefore there is abundant opportunity for variation in the host ability to suppress a TE and the ability of the TE to transpose.

Since the discovery of the piRNA repression system for TEs, the lifecycle of a TE in a host has been envisioned as three steps. First, the TE invades a novel population or species and amplifies unencumbered. TE proliferation is then slowed by segregating insertions in piRNA clusters, and finally inactivated by fixation of piRNA cluster insertions (Kofler 2019). However, clearly bursts, or activity, continues at some level within the population as many of the potentially active TEs in *D. serrata* have a high SFS. This indicates that the TEs have been in the population long enough to accumulate SNPs, potentially including copies with different SNPs continuing to proliferate in the population. It is true that suppression by piRNA cluster insertion may be unstable, but exactly why that is or how important it is for TE survival is not clear.

The accumulation of TEs in laboratory lines should be associated with fitness declines, and be eliminated by selection (Nuzhdin et al. 1997). However, accumulation of TE insertions in individual genotypes, or overall, in genotypes kept in small mass cultures appears to be the rule rather than the exception (Pasyukova 2004; Rahman et al. 2015; Signor 2020). Muller’s rachet may be responsible for the accumulation of insertions, even if they are deleterious (Muller 1932; 1964). What is clear is that TEs are important sources of spontaneous mutations in *Drosophila*, and that in laboratory lines, over time, they may make up a large fraction of the total number of mutations in particular genotypes.

## Availability of data and materials

All raw data is available at NCBI SRA PRJNA410238.

## Acknowledgements

S.S. would like to thank C. & S. Emery for insightful commentary on the manuscript.

## Authors’ contributions

S.S. conceived of the study, performed bioinformatics, and drafted portions of the manuscript. Z.T. performed bioinformatics, interpreted the data, and contributed to the manuscript draft.

